# Dynamic molecular architecture of the synaptonemal complex

**DOI:** 10.1101/2020.02.16.947804

**Authors:** Simone Köhler, Michal Wojcik, Ke Xu, Abby F. Dernburg

## Abstract

During meiosis, pairing between homologous chromosomes is stabilized by the assembly of a protein lattice known as the synaptonemal complex (SC). The SC ensures the formation of crossovers between homologous chromosomes and also regulates their distribution. However, how the SC regulates crossover formation remains elusive. We isolated an unusual mutation in *C. elegans* that disrupts crossover interference but not the assembly of the SC. This mutation alters the unique C-terminal domain of an essential SC protein, SYP-4, a likely ortholog of the vertebrate SC protein SIX6OS1. To characterize the structure of the SC in wild-type and mutant animals, we use three-dimensional STochastic Optical Reconstruction Microscopy (3D-STORM) to interrogate the molecular architecture of the SC in intact germline tissue from *C. elegans*. The approach enabled us to define positions of protein epitopes with respect to the 3D architecture of this complex. Using a probabilistic mapping approach to analyze super-resolution image data, we detect a marked structural transition in wild-type animals that coincides with crossover designation. We also found that our *syp-4* mutant subtly perturbs SC architecture. Our findings add to growing evidence that the SC is an active material whose molecular organization contributes to chromosome-wide crossover regulation.

## Introduction

Diploid genomes are partitioned during meiosis to produce haploid daughter cells. To this end, homologous chromosomes must pair and undergo crossover recombination. This process creates recombinant chromatids that give rise to genetically diverse progeny, and also generates physical linkages that direct proper chromosome segregation.

Programmed double-strand breaks (DSBs) are induced during early meiotic prophase, but only a small subset of DSBs eventually become crossover (CO) recombination events (1). In most eukaryotes, crossovers are nonrandomly far apart, often occurring only once per chromosome pair per meiosis, while other breaks are repaired to yield non-crossover products. It is still unclear how this regulation of crossover number and distribution is achieved.

The synaptonemal complex (SC) plays essential roles in crossover formation and patterning. This protein structure assembles between homologous chromosomes during meiotic prophase, mediating and stabilizing their parallel alignment (2). In transmission electron micrographs the SC appears as a periodic, ladder-like structure when stained with heavy metals (3). Prior work in several organisms has shown that extended coiled-coil proteins, often referred to as “transverse filaments,” span the distance between paired chromosome axes in a head-to-head (N-terminus to N-terminus) configuration, while other proteins localize to the more electron-dark region in the center, known as the “central element.” *In vivo* imaging has revealed that this ordered material shows dynamic, liquid-like behavior despite its highly regular appearance (4-8).

SC proteins self-assemble through a regulated initiation process to form a regular array of proteins that holds paired chromosome axes at a regular distance. These proteins contain extensive regions of potential coiled-coils, some of which presumably mediate specific interactions with each other (9). SC proteins must also interact with proteins associated with the chromosome axes, since mutations in some axis components disrupt assembly of SCs between chromosomes and can lead to self-assembly of “polycomplexes,” dynamic 3D bodies that contain internal striations that recapitulate the dimensions and organization of SCs (5, 10, 11). Some SC proteins also contain peptide motifs that recruit proteins that contribute to crossover patterning and regulate SC assembly and disassembly, including enzymes that mediate posttranslational modifications such as SUMOylation and phosphorylation (8, 12, 13). Very few of these self-assembly and regulatory interfaces within the SC have been identified or characterized in detail. A molecular understanding of the roles and regulation of the SC has been limited in part by a lack of structural information about the physical disposition of such functional elements within the SC.

Six structural proteins essential for SC assembly in *C. elegans* have been identified and named SYP-1–6. SYP-1–4 are each unique, while SYP-5 and SYP-6 are paralogs resulting from a recent gene duplication (9, 14-18). Reduced expression of some SC proteins or mutations that impair or abolish synapsis can perturb CO regulation (18-25). Some of the key proteins required for CO designation localize along the SC prior to their accumulation at eventual CO sites (5, 22, 26-30). It has therefore been proposed that diffusion of CO factors along the SC may govern CO patterning along the length of the chromosomes (29, 31, 32). This idea has been supported by direct visualization of diffusion of meiotic RING finger proteins along the SC (31, 33).

The length and/or fluorescence intensity of the SC can change during the pachytene stage (4, 6, 20). In *C. elegans*, SC proteins also appear to become more stably associated with this structure following the appearance of six bright COSA-1 foci per nucleus, which is indicative of crossover designation. However, it is unclear whether this accumulation is involved in CO patterning or occurs downstream of this process (4, 6-8, 20).

Based on serial-section electron microscopy in various organisms, the central region of the SC is approximately 100 nm in diameter and less than 100 nm in thickness. These dimensions are below the resolution limit of conventional light microscopy. Protein organization within the SC has previously been interrogated through physical interaction assays and immunoelectron microscopy (9, 34), but these methods have limited ability to reveal ultrastructure. Super-resolution fluorescence microscopy has emerged as a powerful tool to investigate the organization of macromolecular assemblies, including the SC (18, 35, 36). In previous work, we established methods to map proteins associated with synapsed chromosome axes using single molecule localization microscopy in intact germline tissue from *C. elegans* (37). Here, we use a similar imaging method to define the organization of individual proteins within the SC central region. In some cases, we mapped two epitopes on the same protein separately, and used a probabilistic method to define the orientation of these proteins based on the distributions of the epitopes. Our findings reveal that the organization of proteins within the SC undergoes a structural transition during pachytene. In the course of this work, we also identified a unique separation-of-function mutation in an unusual SC protein, SYP-4, which disrupts crossover interference and also alters SC ultrastructure. This work expands our understanding of the dynamic organization of the SC during meiosis and its role in crossover patterning.

## Materials and Methods

### Reagents

| **Reagent type** | **Reagent** | **Source** | **Catalog number** |
| --- | --- | --- | --- |
| Primary Antibody | Mouse anti-HA, clone 2–2.2.14 | Thermo Fisher Scientific | 26183 |
| Primary Antibody | Mouse anti-GFP | Sigma Aldrich/Roche | 11814460001 |
| Primary Antibody | Mouse anti-Flag | Millipore Sigma | F1804 |
| Primary Antibody | Rabbit anti-HA | Thermo Fisher Scientific | PA1-985 |
| Primary Antibody | Mouse anti-tubulin | Millipore Sigma | 05-829 |
| Primary Antibody | Goat anti-SYP-1 | Harper et al, 2011 (PMID: 22018922) | N/A |
| Primary Antibody | Rabbit anti-SYP-2 | Colaiacono et al, 2003 (PMID: 12967565) | N/A |
| Primary Antibody | Rabbit anti-pHIM-8/ZIMs | Kim et al, 2015 (PMID: 26506311) | N/A |
| Primary Antibody | Rabbit anti-RAD-51 | Harper et al, 2011 (PMID: 22018922) | N/A |
| Primary Antibody | Chicken anti-HTP-3 | MacQueen et al, 2005 (PMID: 16360034) | N/A |
| Primary Antibody | Rabbit anti-HIM-3 | ModENCODE/SDIX | SDQ4713 |
| Secondary Antibody | Alexa Fluor 647 Donkey anti-mouse | Jackson Immuno Research | 715-605-151 |
| Secondary Antibody | Alexa Fluor 647 Donkey anti-goat | Jackson Immuno Research | 705-605-147 |
| Secondary Antibody | Alexa Fluor 647 Donkey anti-rabbit | Jackson Immuno Research | 711-605-152 |
| Secondary Antibody | Cy3 Donkey anti-mouse | Jackson Immuno Research | 715-165-151 |
| Secondary Antibody | Cy3 Donkey anti-rabbit | Jackson Immuno Research | 711-165-152 |
| Secondary Antibody | Cy3 Donkey anti-chicken | Jackson Immuno Research | 703-165-155 |
| Secondary Antibody | Cy5 Donkey anti-mouse | Jackson Immuno Research | 715-175-151 |
| Secondary Antibody | Cy5 Donkey anti-chicken | Jackson Immuno Research | 703-175-155 |
| Secondary Antibody | Alexa Fluor 488 Donkey anti-mouse | Jackson Immuno Research | 715-545-151 |
| Secondary Antibody | Alexa Fluor 488 Donkey anti-chicken | Jackson Immuno Research | 703-545-155 |
| Secondary Antibody | Alexa Fluor 488 Donkey anti-goat | Jackson Immuno Research | 705-545-147 |
| Secondary Antibody | HRP Donkey anti-mouse | Jackson Immuno Research | 715-035-151 |
| Secondary Antibody | Donkey anti-rabbit | Jackson Immuno Research | 711-005-152 |
| Nanobody | Alexa Fluor 488 GFP-booster | ChromoTek | gb2AF488 |
| Fluorophor | CF568 succinimidyl ester | Biotium | NC1542764 |
| Mounting Medium | ProLong Gold antifade | Thermo Fisher Scientific | P10144 |
| CRISPR reagent | CRISPR tracrRNA | Integrated DNA Technologies, Skokie, IL | 1072534 |
| CRISPR reagent | *S. pyogenes* Cas9-NLS | QB3 MacroLab at UC Berkeley | N/A |
| CRISPR reagent | crRNA, repair template, oligos | Integrated DNA Technologies, Skokie, IL | see Table S3 for sequences |
| Western Blotting Detection | ECL Select Western Blotting Detection Reagent | Millipore Sigma | Cytiva RPN2235 |

### Biological Resources

A complete list of *C. elegans* strains used in this study can be found in Table S1. All strains were cultured at 20 ºC using standard methods (38). GFP::SYP-3 is a single copy transgene inserted by MosSCI (39). Other tags were inserted using CRISPR-Cas9 genome editing as described in (37). Briefly, Cas9 and gRNA were delivered by microinjection, either encoded on a plasmid (40) or as *in vitro* preassembled Cas9-ribonucleoprotein complexes. Repair templates for small epitope tags were codon optimized for *C. elegans* (41) and synthesized as “Ultramers” by IDT.

### Immunofluorescence

Gonads were dissected from young adults (24 h post-L4) and immunofluorescence was performed as described (42) with modifications described in (37). The following primary antibodies, all of which have been previously described, were used: goat anti-SYP-1 (1:500, affinity purified) (43), mouse anti-HA (1:500, monoclonal 2-2.2.14, Thermo Fisher Scientific), rabbit anti-SYP-2 (1:500, affinity purified) (15), mouse anti-GFP (1:500, monoclonal 7.1 and 13.1, Roche), mouse anti-Flag (1:500, monoclonal M2, Sigma-Aldrich), chicken anti-HTP-3 (1:500) (44), rabbit anti-HIM-3 (1:500, SDQ4713 modENCODE project) (45), rabbit anti-HIM-8pT64 (1:2000) (46), and rabbit anti-RAD-51 (1:500) (43). Commercial secondary antibodies were raised in donkeys and fluorescently labelled with Alexa Fluor® 488, Cy3 or Alexa Fluor® 647 (1:500, Jackson ImmunoResearch and Invitrogen). Gonads were mounted in ProLong Gold antifade mountant (ThermoFisher Scientific) and epifluorescence images were acquired on a DeltaVision Elite microscope (Applied Precision) using a 100x N.A. 1.4 oil-immersion objective.

To quantify protein levels by immunofluorescence, imaging was performed on a Marianas spinning-disk confocal microscope (3i) with a 100x 1.46 NA oil immersion objective. 3D stacks of wild-type (*syp-4::HA(ie29)*) and *syp-4(ie25)* animals were acquired from the same slide. SC-containing voxels were identified and quantified in Fiji by thresholding the SYP-2 and SYP-4::3xFLAG channels for *syp-4(ie29)* and *syp-4(ie25)*, respectively. The intensities of segmented SCs were then analyzed using the “3D Object Counter” plugin. Images shown in Supplementary Fig. S5 are background subtracted maximal intensity projections with equally scaled intensity values.

### STORM and PALM imaging

Super-resolution imaging of immunostained, intact germline tissue was carried out as described (37). For STORM, secondary antibodies raised in donkeys or goats labelled with Alexa Fluor® 647 were used (1:500, Jackson ImmunoResearch and Invitrogen). Following acquisition of the antibody signal, the fluorescently tagged internal reference protein, mEos2::HIM-3 or mMaple3::HIM-3 (37) was imaged using PALM (47, 48). For early pachytene images, mEos2::HIM-3 was co-stained with a rabbit anti-HIM-3 antibody and a donkey-anti-rabbit secondary antibody (Jackson ImmunoResearch) labelled with CF568-NHS ester to achieve a 2:1 dye-to-antibody ratio. All images were rotated about the optical axis such that the direction of the chromosome axes defines the *y* axis. Aligned and averaged images (Supplementary Fig. S3) were then used to generate histograms of localization events in *x* and *z* (37). To systematically distinguish between mono- and bi-modal distributions, we evaluated fits with one and two Gaussians using an ANOVA test in R (*p* < 0.05). Standard deviations of fit parameters were estimated by a subsampling approach using subsets of half the number of individual SC stretches (37). The results are summarized in Table S2.

### Statistical analysis

Sample sizes were not predetermined, and experiments were not randomized. The investigators were not blinded to allocation during experiments and outcome assessment. For STORM experiments, the total lengths and number of stretches analyzed are summarized in Table S2. p-Values were calculated by two-sided Mann-Whitney-Wilcoxon tests in R (version 4.1.2).

### Analysis of SC organization

To derive a model of SC organization from our data, we treated each protein as a rigid rod and determined which orientation was most consistent with the distributions of localization events mapped by STORM. We first removed the most extreme 2.5% localization events using squared Mahalanobis distances in R (version 3.4.2). We then mapped each localization event corresponding to the *n*th-percentile in *x* and the *m*th-percentile in *z* of the N-(or C-) terminal distribution to a randomly selected localization event within the *n*th ± 7.5% in *x* and *m*th ± 7.5% in *z* of the corresponding C-(or N-) terminal distribution. This analysis yields a distance between each pair of N- and C-terminal localization events and an orientation of the protein within the SC. Each localization event may be as far as 20 nm from the epitope, since we used unlabeled primary and dye-labeled secondary antibodies; thus, we expect that these distances may vary by ±40 nm relative to the true distance. Some pairs of localization events between the N- and C-terminal distributions yield improbably short or long distances between N- and C-terminal antibodies. We restricted our analysis to mapping events that yield distances between the 5th and 95th percentile of all distances mapped. Using this approach, the distances for all conditions were between 11.6 nm and 120 nm (Figure S4B).

### Recombination mapping

To map meiotic crossovers in *syp-4*^*CmutFlag*^*(ie25)* homozygotes, this allele (which was generated in the N2 Bristol background) was introgressed into the divergent Hawaiian strain CB4856 by 8 sequential crosses. Hawaiian and N2 strains homozygous for either wild-type *syp-4* or *syp-4*^*CmutFlag*^*(ie25)* were crossed to each other, and the hybrid F1 progeny were backcrossed to Hawaiian *syp-4*^*+*^ males. The resulting F2 progeny inherit one haploid complement of CB4856 alleles and a set of potentially recombinant N2/CB4856 maternal chromosomes. To map recombination events, F2 hermaphrodites were plated individually, allowed to reproduce for one generation, and the genomic DNA of their pooled progeny was isolated by phenol-chloroform extraction. Illumina sequencing libraries were prepared as described in (50) and sequenced as 50 bp single reads on a HiSeq2500 Illumina sequencer at the Vincent J. Coates Genomics Sequencing Laboratory at UC Berkeley, with a coverage of 2-4x genomes per sample. Reads were mapped to reference genomes of the N2 (Wormbase release WS230) and CB4856 (51) strains. Genotypes were called using the multiplexed shotgun genotyping (MSG) toolbox with default parameters (52). The aligned sequences are available through the NCBI SRA database under accession number SRP126693. Crossovers that occurred in oogenesis were detected as transitions from Hawaiian/Bristol heterozygous stretches to homozygous Hawaiian stretches. The observed number of crossovers was doubled, since only half of the total that occurred in each meiotic cell are inherited by individual progeny.

### Electron microscopy

High-pressure freezing, freeze-substitution, sample preparation, and microscopy were performed as described previously (5, 53, 54). Images were acquired on a Tecnai 12 transmission electron microscope (120 kV, FEI, Hillsboro, OR) equipped with a Gatan Ultrascan 1000 CCD camera (Pleasanton, CA).

### Immunoblotting

To compare expression levels of SYP proteins, 120 adult worms were lysed by boiling in 40 µL Laemmli sample buffer with β-mercaptoethanol for about 5 min, until particulate matter was not detected using a dissection stereomicroscope. Whole-worm lysates were electrophoresed using Nupage 4-12% polyacrylamide gradient gels and transferred to PVDF membranes. Primary antibodies were rabbit (Pierce, PA1-985, used for HA::SYP-1) or mouse anti-HA (SYP-4) and mouse anti-α-tubulin (DM1A, EMD Millipore), each diluted 1:5,000. HRP-conjugated secondary antibodies (Jackson Laboratory) were detected using ECL reagents (Amersham). SYP-4::HA and tubulin were detected using the same HRP-conjugated anti-mouse 2° antibody. HA::SYP-1was detected using HRP-anti-rabbit/ECL and tubulin was detected by Cy3-conjugated anti-mouse secondary antibody (1:5,000). Images were recorded using a Chemidoc system (Bio-Rad) and quantified using Fiji.

## Results

### A separation-of-function mutation in the C-terminus of SYP-4 perturbs crossover interference

To map protein epitopes using STORM super-resolution microscopy, we inserted epitope tags into the endogenous genes encoding the synaptonemal complex (SC) proteins SYP-1, SYP-2, SYP-3, and SYP-4 in *C. elegans* using CRISPR/Cas9 genome editing.

All tagged alleles were characterized through brood counts (Supplementary Table S1). Production of inviable embryos and male self-progeny are indicative of chromosome mis-segregation during meiosis (55). Broods from wild-type hermaphrodites show embryo viability close to 100%, with approximately 0.2% male self-progeny.

Fortuitously, we obtained three different C-terminally tagged alleles of SYP-4 that had distinct effects on meiosis (Fig. 1A,B). Insertion of an HA tag at the C-terminus (*syp-4*^*ha*^) did not disrupt its function, as indicated by brood analysis (Fig. 1C). An independent insertion of HA at the C-terminus also had a noncoding insertion within the 3’UTR (*syp-4*^*ha3’UTR*^), which reduced protein expression to about 30% of that seen in *syp-4*^*ha*^ (Fig. 1B). A third allele, designated as *syp-4*^*CmutFlag*^, incorporated a 3xFLAG tag at the C-terminus and also had an altered C-terminal sequence due to a frame-shift mutation. The *syp-4*^*ha3’UTR*^ and *syp-4*^*CmutFlag*^ alleles partially compromised the function of SYP-4, which was initially revealed by reduced progeny viability and elevated male self-progeny (Fig. 1C, Supplementary Table S1).

**Figure 1:**
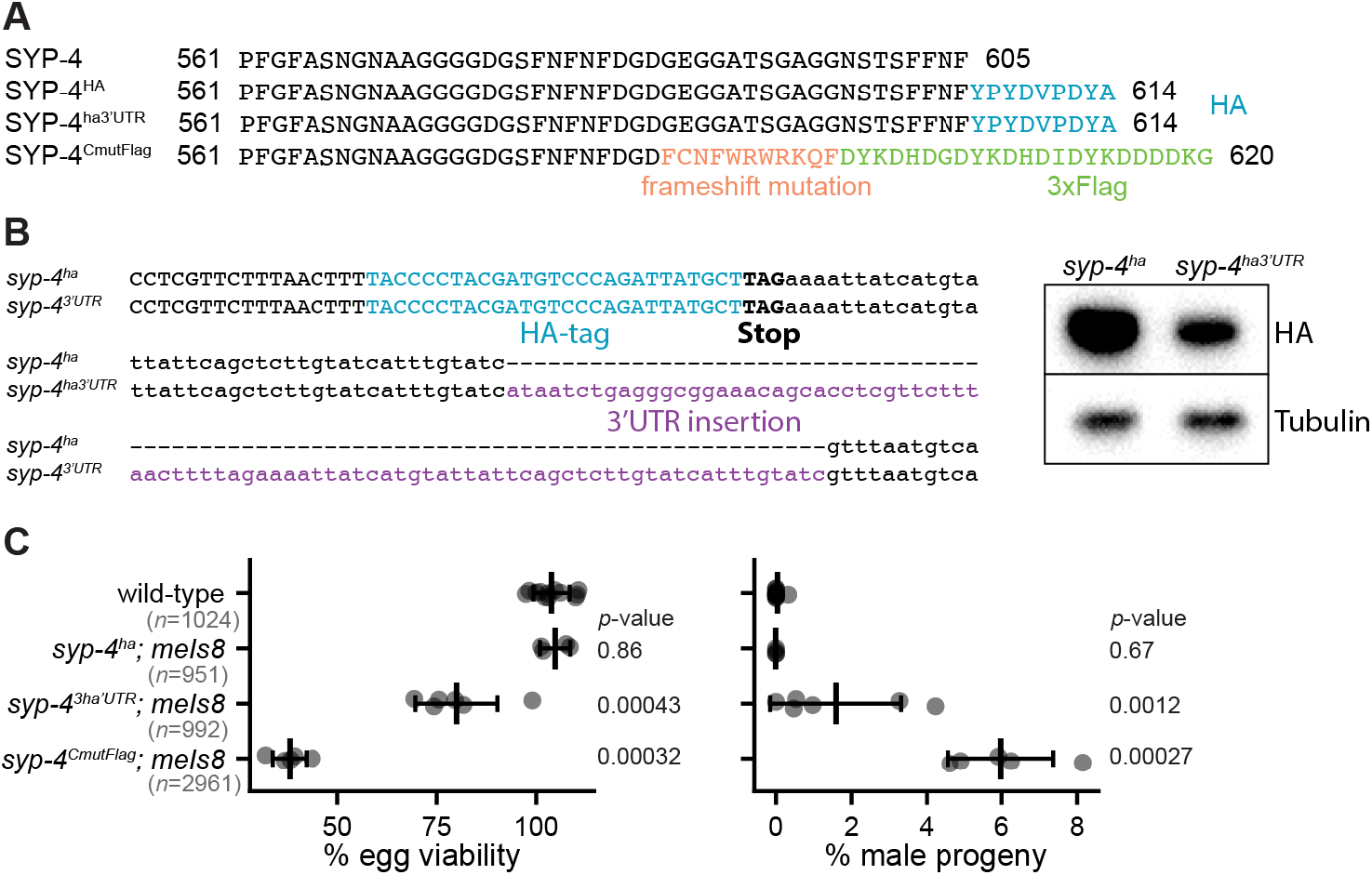
Identification of two partially functional SYP-4 alleles. (A) The C-terminal amino-acid sequences of wild-type SYP-4, SYP-4^ha^, SYP-4^ha3’UTR^ and SYP-4^CmutFlag^ are indicated. HA-tags are highlighted in blue, mutated residues in orange, and 3xFlag peptide in green. (B) Genome sequences for *syp-4*^*ha*^ and *syp-4*^*ha3’UTR*^ are shown (left). An insertion within the 3’UTR (purple) of *syp-4*^*ha3’UTR*^ caused a reduction in protein expression compared to *syp-4*^*ha*^, as revealed by western blotting (right). (C) Egg viability is reduced and the incidence of male progeny is increased in *syp-4*^*ha3’UTR*^ and *syp-4*^*CmutFlag*^ mutant animals, while *syp-4*^*ha*^ animals resemble wild-type animals. Black bars show mean±s.d. values, dots show values for individual animals scored, *n* denotes the total number of progeny scored.

Immunofluorescence revealed continuous SC along the entire lengths of the chromosome axes in HA-tagged *syp-4*^*ha*^ animals (Fig. 2A). Some partial and delayed SC assembly was seen in *syp-4*^*ha3’UTR*^ animals, consistent with evidence that reduced expression of SYP-1, -2, or -3 can perturb synapsis (Figure 2B, (20)). Unexpectedly, however, SC assembly appeared to be complete and timely in *syp-4*^*CmutFlag*^ animals (Figure 2C), despite the high nondisjunction they displayed. A fluorescent reporter for CHK-2 kinase activity(46) revealed a delay in meiotic progression in both *syp-4*^*CmutFlag*^ and *syp-4*^*ha3’UTR*^ animals (Fig. 2D). CHK-2 promotes pairing, synapsis, and double-strand break (DSB) induction; it is normally active in early meiotic prophase and then decays at mid-pachytene, concomitant with crossover designation, but its activity is prolonged by a “crossover assurance checkpoint” under conditions that impair synapsis and/or crossover designation (46) (56). Consistent with prolonged CHK-2 activity, we also observed an extended region of nuclei positive for RAD-51 foci, and more foci per nucleus, in both *syp-4*^*CmutFlag*^ and *syp-4*^*ha3’UTR*^ compared to wild-type animals. (Supplementary Fig. S1). This may reflect a delay in establishment of crossover intermediates in these mutants, or a defect in cells’ ability to detect their presence and inactivate the crossover assurance checkpoint. Importantly, six bivalents were consistently observed at diakinesis in all three tagged *syp-4* strains, indicating that crossovers eventually occurred on all chromosomes (Fig. 2E). However, the structure of bivalents at diakinesis was altered in *syp-4*^*CmutFlag*^ oocytes (Supplementary Fig. S2), indicating that crossover number might be perturbed (57-59).

**Figure 2:**
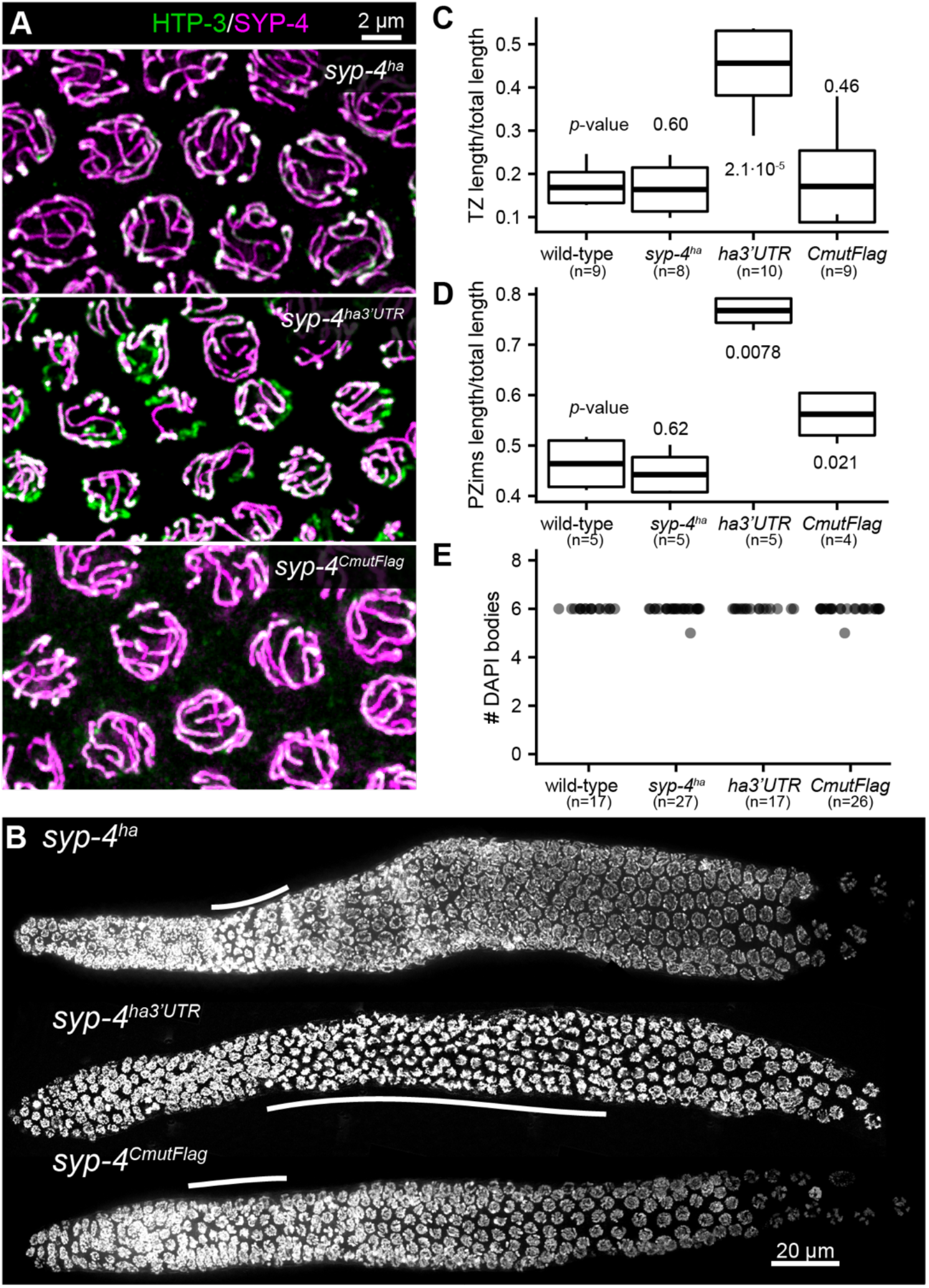
Synapsis is delayed in SYP-4 depleted *syp-4*^*ha3’UTR*^ but not in *syp-4*^*CmutFlag*^ mutant animals. (A) Diffraction-limited images reveal robust synapsis for *syp-4*^*ha*^ (top) and *syp-4*^*CmutFlag*^ (bottom) mutant animals with overlapping HTP-3 (green) and SYP-4 (magenta) staining in mid pachytene. By contrast, some unsynapsed axes (green stretches) are observed at mid-pachytene in *syp-4*^*ha3’UTR*^ animals (middle). (B) Elongation of the transition zone, as defined by nuclei with crescent-shaped DAPI-bright regions, is observed in *syp-4*^*ha3’UTR*^ animals but not in *syp-4*^*ha*^ or *syp-4*^*CmutFlag*^ animals, consistent with the defects in synapsis. Quantification of transition zone lengths is shown in (C). (D) CHK-2 activity is prolonged in *syp-4*^*CmutFlag*^ mutants. In wild-type or *syp-4*^*ha*^, the “CHK-2 active” zone detected with an antibody against a CHK-2-dependent phosphoepitope on pairing center proteins (Kim 2015) comprises 45% of the length of the region spanning the leptotene, zygotene and pachytene stages of meiosis. This fraction was extended to 77% in *syp-4*^*ha3’UTR*^, and to 56% in *syp-4*^*CmutFlag*^ animals. Boxplots show mean±s.d. (whiskers show minima and maxima). (E) Despite varying delays in meiotic progression, oocytes homozygous for all *syp-4* alleles show 6 DAPI staining bodies at diakinesis, indicative of crossover formation on all chromosomes.

To analyze crossover formation in our tagged *syp-4* strains, we crossed in GFP::COSA-1, which marks designated crossover sites at late pachytene (60). Wild-type nuclei display 6 bright GFP::COSA-1 foci in late pachytene, corresponding to a single CO site per chromosome (57, 60). Knockdown of SC proteins can elevate this number to 7-9 per nucleus (20). Consistent with these previous findings, we observed 6±0.2 (s.d.) GFP::COSA-1 foci in wild-type oocytes and 7.3±1.1 foci in *syp-4*^*3’UTR*^. Remarkably, we observed 10.9±1.7 GFP::COSA-1 foci at late pachytene in *syp-4*^*CmutFlag*^ hermaphrodites (Fig. 3A,B), surpassing the numbers previously observed in any other situation. We also measured crossing-over genetically using whole genome sequencing (see methods). This confirmed that the increase in GFP::COSA-1 foci in *syp-4*^*CmutFlag*^ animals quantitatively reflects an increase in COs (Fig. 3B).

**Figure 3:**
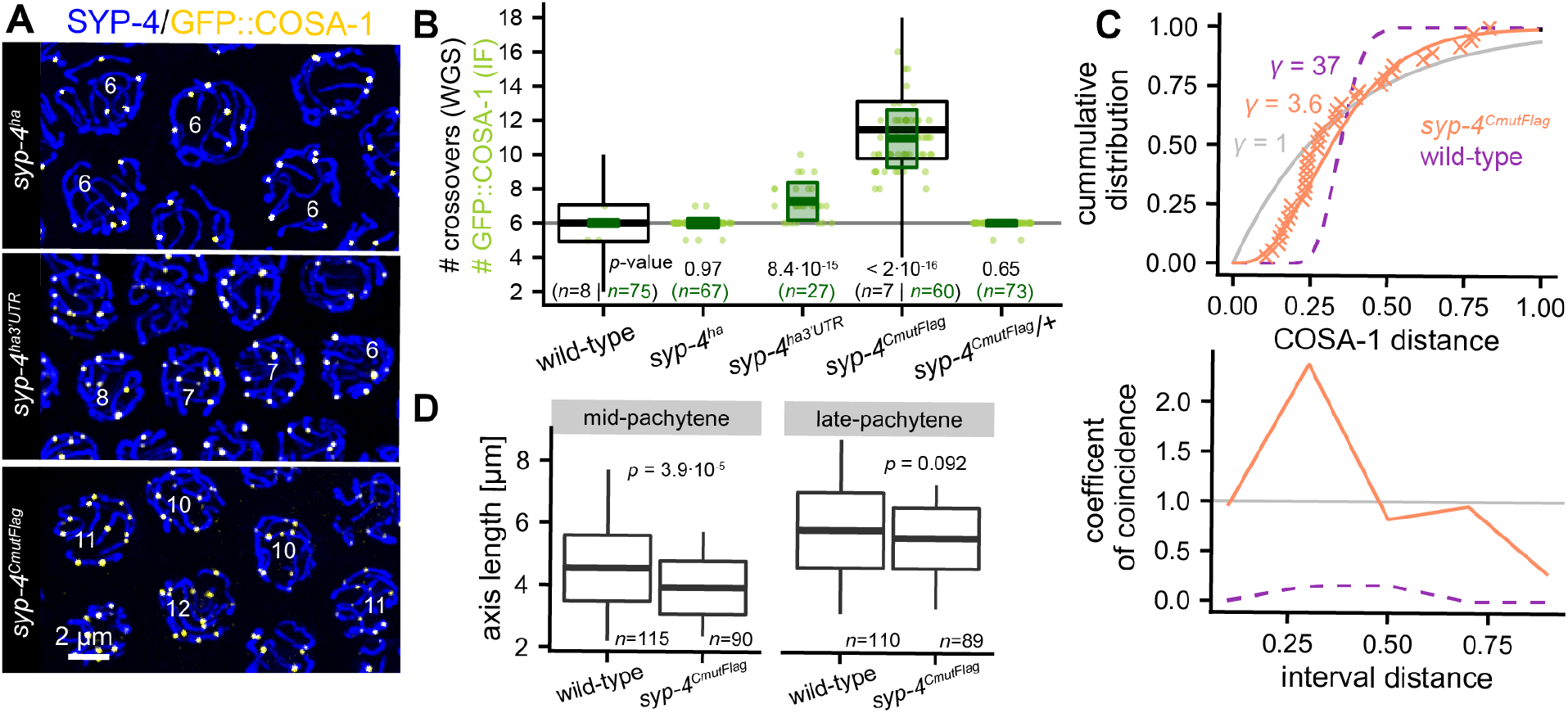
Crossover interference is abrogated in *syp-4*^*CmutFlag*^ mutants. (A) Maximum-intensity projections of deconvolved 3D widefield images of late pachytene nuclei stained for GFP::COSA-1 (yellow), which marks designated crossover sites, and SYP-4 (SC) (blue). (B) Correlation between designated crossover sites detected as GFP::COSA-1 foci and crossovers measured through whole-genome sequencing (WGS) of progeny of hybrid animals. Cytological data are shown as green dots, with boxes to indicate mean±s.d.; green numbers below indicate the number of nuclei scored. Crossover numbers based on WGS for wild-type and *syp-4*^*CmutFlag*^ (mean±s.e.; whiskers show extreme values), are overlaid in black. *syp-4*^*ha3’UTR*^ and *syp-4*^*CmutFlag*^ mutant animals show elevated numbers of GFP::COSA-1 foci, while *syp-4*^*ha*^ and heterozygous *syp-4*^*CmutFlag*^*/+* hermaphrodites are indistinguishable from wild-type. (C) To quantify crossover interference strength in *syp-4*^*CmutFlag*^ mutant animals, we fit the cumulative distribution function of inter-GFP::COSA-1 distances in *syp-4*^*CmutFlag*^ mutants (orange crosses) using a gamma function (top panel) (61). *Γ*=1 denotes no interference (gray), *γ*=37 corresponds to interference measured for wild-type animals (20) (purple dashed line), while interference in *syp-4*^*CmutFlag*^ homozygotes is severely reduced to *γ*=3.5 (orange line, *n*=41 chromosomes). Alternatively, interference strength can be described by the coefficient of coincidence (bottom panel) (62), which denotes the ratio of the observed number of 2 COs (GFP::COSA-1 foci) at a given distance to the expected number assuming random CO distribution. In wild-type animals, the coefficient of coincidence of GFP::COSA-1 foci is 0, corresponding to complete interference (purple dashed line; *n*=88 chromosomes), while it is approximately 1 in *syp-4*^*CmutFlag*^ mutants (orange, *n*=41), indicating an absence of interference (gray line). (D) Chromosome axes lengths are shorter in *syp-4*^*CmutFlag*^ mutants compared to wild-type axes in mid-pachytene prior to designation of CO sites (before occurrence of bright GFP::COSA-1 foci, left) and comparable in late pachytene (right). Boxplots show mean±s.d and whiskers are extreme values.

We quantified crossover interference in *syp-4*^*CmutFlag*^ using two different metrics: the gamma factor (61) and the coefficient of coincidence (62). By either measure, crossover interference was severely reduced or absent in *syp-4*^*CmutFlag*^ oocytes (Fig. 3C). The increase in crossing-over is sufficient to explain the high nondisjunction we observed, since excess crossovers interfere with proper chromosome segregation (59). Crossover interference is a distance-dependent effect, and in many organisms longer chromosomes undergo more crossovers than shorter ones (63-66). Surprisingly, we found that chromosome axes in *syp-4*^*CmutFlag*^ were shorter than in wild-type animals (Fig 3D). Thus, changes in chromosome length cannot account for the observed increase in COs in *syp-4(ie25)* animals.

### Loss of ‘interference’ in polycomplexes with C-terminally mutated SYP-4

We also examined the effect of *syp-4*^*CmutFlag*^ in the context of polycomplexes. These are ordered assemblies of SC proteins that form in the nucleoplasm when SC cannot assemble between chromosomes. In wild-type *C. elegans* oocytes, polycomplexes are often observed transiently prior to synapsis, and larger polycomplexes are observed throughout prophase in mutants lacking the axis protein HTP-3 (5, 10, 11) (Fig. 4A). Despite an absence of double-strand breaks, polycomplexes in *htp-3* mutants recruit several pro-CO proteins, which undergo dynamic redistribution during meiotic progression (5). In particular, the RING finger proteins ZHP-3 and ZHP-4 are initially distributed throughout polycomplexes, but later concentrate at foci that are also positive for COSA-1, and thus resemble the “recombination nodules” that normally mark designated CO sites along SCs in late pachytene (29). In *htp-3* mutant animals that express wild-type SC proteins, most polycomplexes show a single focus of these pro-crossover proteins at mid-prophase (1.1±0.2; Fig. 4A,D). Intriguingly, in *htp-3 syp-4*^*CmutFlag*^ double mutants, we frequently observed multiple GFP::COSA-1 foci (2.1±0.8) associated with individual polycomplexes (Fig. 4C,D), mirroring the ∼2-fold increase in GFP::COSA-1 foci observed along bona fide SCs in *syp-4*^*CmutFlag*^ mutants. Thus, the observed increase in the number of GFP::COSA-1 foci in *syp-4*^*CmutFlag*^ mutant animals was recapitulated in polycomplexes devoid of CO precursors. By contrast, when we combined *htp-3* and *syp-4*^*3’UTR*^, we observed small polycomplexes associated with a single (1.1±0.2) COSA-1 focus (Fig. 4B,D).

**Figure 4:**
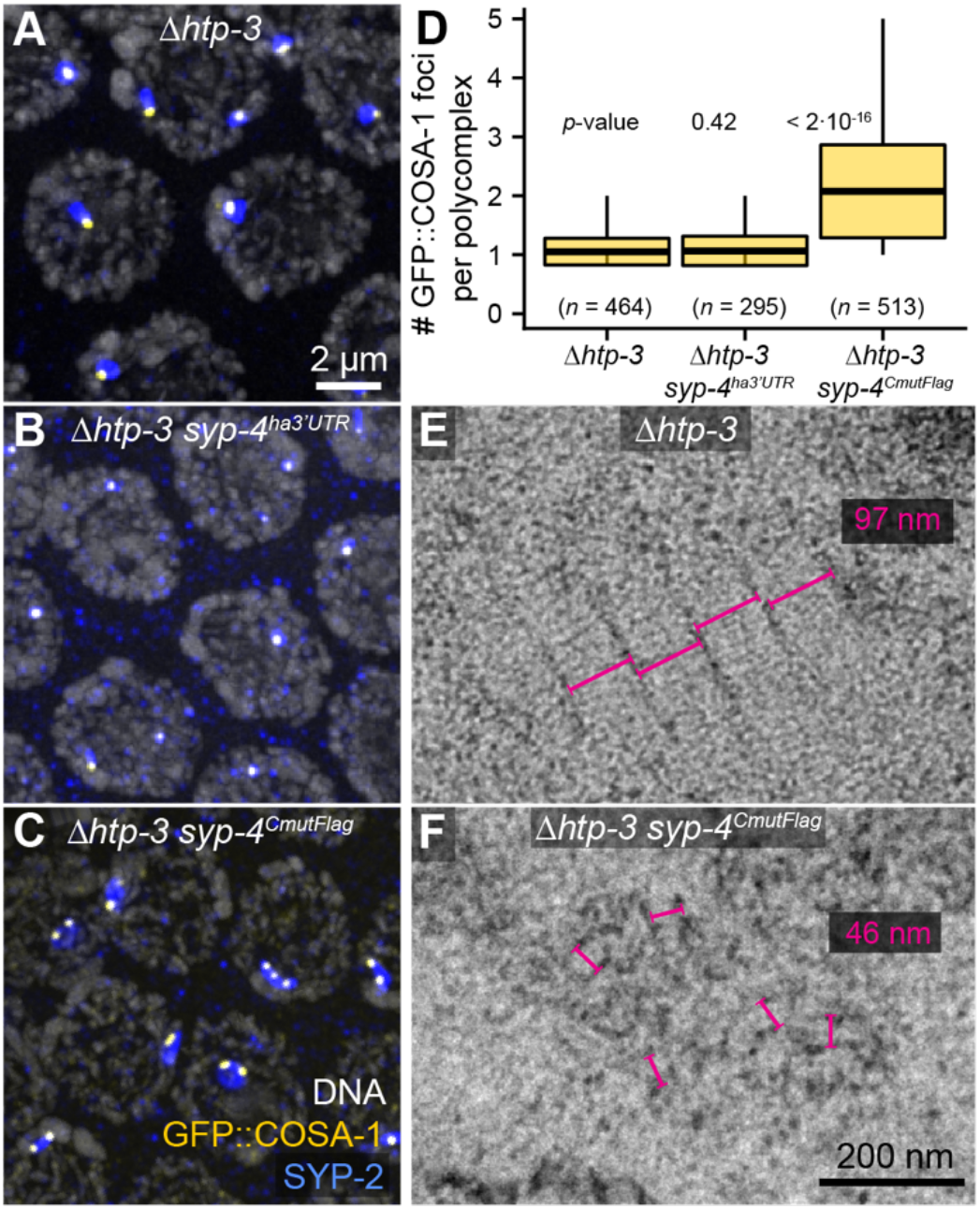
Irregular organization and an increase in GFP::COSA-1 foci in *syp-4*^*CmutFlag*^ polycomplexes. (A-C) Fluorescence micrographs show that the number of GFP::COSA-1 foci (yellow) is limited to 1 or 2 per polycomplex (SYP-2, blue) in *htp-3* animals expressing wild-type SC proteins (A) or SYP-4^HA3’UTR^ (B) but is elevated in *htp-3 syp-4*^*CmutFlag*^ double mutants (C). DAPI is shown in gray. (D) Quantification of GFP::COSA-1 foci. Boxplots show mean±s.d. and whiskers are extreme values. (E, F) Representative electron micrographs show that the SC-like structure of polycomplexes (E, *̄htp-3*, 97-nm spacing of parallel, electron dark regions) is disturbed in *Δhtp-3 syp-4*^*CmutFlag*^ (F), which are disorganized and 46 nm apart. This may reflect a destabilization of the midline region, or “central element,” of the SC within polycomplexes.

We next investigated the organization of polycomplexes using transmission electron microscopy (TEM). As in other organisms, polycomplexes in *C. elegans* recapitulate the periodic organization of SCs and are thus presumed to represent multiple SC units that form a non-isotropic, 3-dimensional liquid crystalline lattice. We observed these characteristic striated complexes in *htp-3* mutant animals (Fig. 4E). By contrast, in *htp-3 syp-4*^*CmutFlag*^ double mutants, we observed nuclear aggregates that resembled polycomplexes, with electron-dense lateral regions interspersed with more electron-lucent central regions, but lacked the clear striated central pattern that is characteristic of polycomplexes in *htp-3* single mutants (Fig. 4E,F). The polycomplexes in *htp-3 syp-4*^*CmutFlag*^ double mutants appeared to be internally fragmented, rather than maintaining regular orientation over hundreds of nanometers (Fig. 4E, (5)), and the distance between parallel electron-dark bands was much narrower (46.3±1.2 nm, vs 97.6±1.5 nm in *htp-3* polycomplexes, Fig. 4E,F). Thus, *syp-4*^*CmutFlag*^ perturbs the organization of proteins within polycomplexes, and also disrupts the distribution of pro-CO factors associated with each of these structures, suggesting a direct link between the ultrastructure of the SC and the regulation of CO patterning.

### Dynamic architecture of the synaptonemal complex in *C. elegans*

While transmission electron microscopy first revealed the ordered, striated structure of the SC and polycomplexes, its ability to define the molecular organization of these assemblies is limited. We therefore took advantage of 3D-STORM super-resolution microscopy (67, 68) to map the orientation of components within the SC, an approach that we have recently used to define the molecular architecture of meiotic chromosome axes and the orientation of two recently discovered components of the SC (18, 37) (Supplementary Fig. S3). To map the relative localization of individual epitopes of proteins in the SC, we imaged the same fluorescently tagged reference protein (HIM-3 tagged with mEos2 or mMaple3, (69)) in all samples using PALM (photoactivated localization microscopy, Supplementary Fig. S3B-C). Regions in which paired axes were in the same focal plane were selected, so that all images were collected parallel to the SC, and could then be co-oriented by rotation about the optical axis. Because paired axes are held at a constant distance when the SC assembles between, we could average data from multiple images to define the distributions and average positions of protein epitopes. We used a 3D coordinate system in which the orientation of the chromosome axes defines the *y*-axis, the *x*-axis is thus orthogonal to both chromosome axes, and *z* the optical axis. The distributions of each protein epitope could then be mapped in *x* and *z*.

To investigate whether the organization of the SC along homologous chromosomes is affected by *syp-4*^*CmutFlag*^, we first analyzed the organization of one of the transverse filament proteins in the *C. elegans* synaptonemal complex, SYP-1 (14, 34). To this end, we localized an antibody raised against the C-terminal peptide sequence of SYP-1 (43). In frontal view, this SYP-1 epitope was resolved as two parallel strands, located at 42.3±0.8 nm and 42.0±1.2 nm off-center in early and late pachytene of wild-type animals, respectively (Fig. 5A). The width of the SC central region in *C. elegans* is ∼96 nm (5); this epitope thus lies near the outer edges of the central region and within a few nanometers of HIM-3, the most proximal known component of the chromosome axis (37) as expected for a transverse filament protein (34).

**Figure 5:**
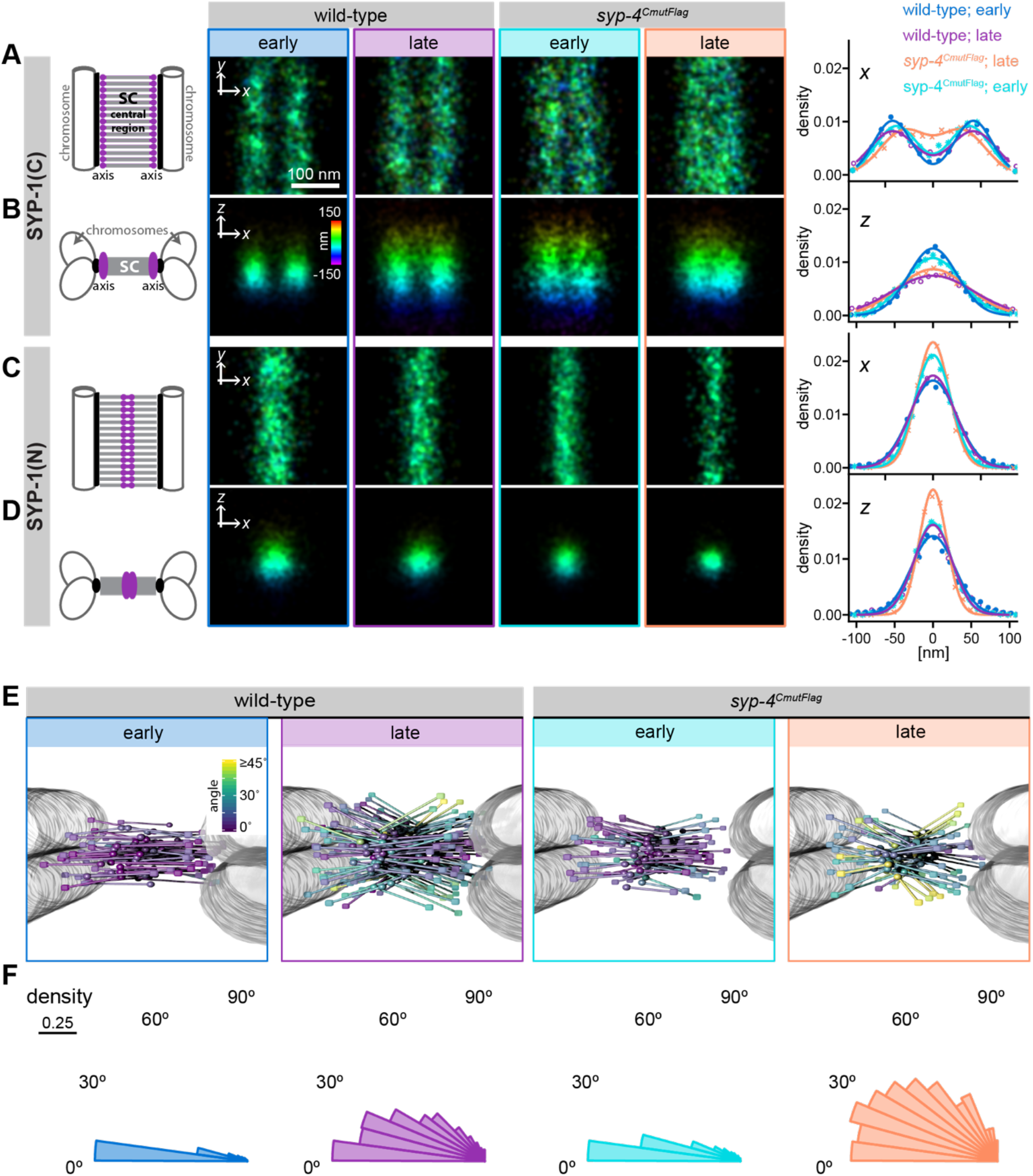
Super-resolution microscopy reveals distinct three-dimensional organizations of SYP-1 during pachytene and in *syp-4*^*CmutFlag*^ mutant animals. (A) Measurements based on averaged STORM images indicate that the C-terminus of SYP-1 lies close to the chromosome axes. Colors denote localizations in *z* from -150 to 150 nm. Synaptonemal complexes in early and late pachytene are shown in frontal view (cartoon, left), and plots (right) show histogram of localization events. Histograms are fit with 1 or 2 Gaussians. The results of the fits are summarized in Supplementary Table S2. (B) Cross-sectional views of 3D-STORM images in (A) are shown. The N-terminus of SYP-1 is localized near the center of the SC both in frontal (C) and cross-sectional (D) view. (E) The stochastic nature of STORM does not allow us to image the N- and C-terminus of a specific single molecule at the same time, but will yield population averages for the localization of individual domains. We therefore used a probabilistic mapping approach to model the orientation of SYP-1 molecules for each condition. The resultant models of SYP-1 are shown with C-termini depicted as cubes and N-termini as spheres. Colors of SYP-1 molecules indicate the angle with respect to the central plane to the SC. A proxy for chromatin (gray areas) is added for visualization purposes. Models are rendered using POV-ray (v3.7.0). Models show changes in the orientation of SYP-1 during pachytene and in *syp-4*^*CmutFlag*^ mutant animals. (F) These changes are revealed by the distributions of angles of SYP-1 with respect to the central plane of the SC in cross-sectional views. Scale bars in (A) and (B) apply to all images.

Two discrete distributions could also be resolved for the SYP-1 C-terminus in *syp-4*^*CmutFlag*^ mutants. However, the inter-peak distance was reduced to 38.0±2.7 nm and 30.5±1.4 nm in early and late pachytene, respectively. This finding is consistent with the closer spacing between electron-dark regions in *htp-3 syp-4*^*CmutFlag*^ compared to *htp-3* polycomplexes (Fig. 4E,F).

We also measured the localization of the SYP-1 C-terminus along the *z*-axis (Fig. 5B). In wild-type animals, we observed that the distribution of localization events in *z* was narrower in early than in late pachytene (*xz*-view, Fig. 5B, Table S2). This is consistent with evidence that SYP-1 may accumulate along the SC later in pachytene (4, 6), which may result in a thicker complex. In contrast, in *syp-4*^*CmutFlag*^ mutant animals, the *z*-distribution of SYP-1 C-termini remained narrower than in wild-type animals in late pachytene (Fig. 5B), suggesting that this mutation may reduce the tendency of the SC to grow in thickness concomitant with crossover designation. This may be a consequence of the dysregulation of meiotic progression in this mutant, as indicated by prolonged CHK-2 activity described above.

We then asked whether the N-terminus of SYP-1 is similarly altered during meiotic progression or in *syp-4*^*CmutFlag*^ mutants. Tagging the N-terminus of SYP-1 disrupts SC assembly ((70) and data not shown), but we identified a poorly conserved region close to the N-terminus which accommodated insertion of an HA epitope without impairing SC function (Supplementary Table S1). Consistent with previous evidence of a transverse orientation for SYP-1 (34), the N-terminus was confined to a single peak at the center of the SC along the *x*-axis in both early and late pachytene in both wild-type and *syp-4*^*CmutFlag*^ mutant animals. However, the N-terminal distribution did not change detectably during pachytene, nor was it altered in the *syp-4*^*CmutFlag*^ strain (Fig. 5C,D). This finding suggested that the increase in thickness we observed for the C-terminus of SYP-1 is not caused by a mere stacking of SYP-1 molecules along the *z*-axis but may involve a more complex re-organization of SYP-1 within the SC.

To better define the orientation of SYP-1 within the SC, we modeled the region spanning the two epitopes as a rigid rod, and mapped the N-terminal localization events to the C-terminal distributions (Supplementary Fig. S4, see methods). This approach indicated that SYP-1 lies nearly parallel to the plane of the SC in early pachytene and becomes more splayed in late pachytene (Fig. 5E,F). This reorganization could result from either a change in orientation or conformation of the protein. These results cannot be explained as a simple consequence of accumulation of additional SC layers in late pachytene. Intriguingly, in *syp-4*^*CmutFlag*^ mutant animals SYP-1 is more diagonally oriented in early pachytene, and is even more splayed in late pachytene (Fig. 5E,F). Thus, the *syp-4*^*CmutFlag*^ allele perturbs both the organization of proteins within polycomplexes and SCs assembled between homologous chromosome axes.

We tested whether the conformation of other SC components also changes during meiotic progression or in the presence of *syp-4*^*CmutFlag*^. We mapped the C-termini of SYP-2 using an antibody raised against its C-terminal sequence (15), the C-terminal HA epitopes on SYP-3::HA and SYP-4::HA, the N-terminus of GFP::SYP-3 (39), and a Flag epitope inserted into the middle of SYP-4. We further mapped the C-termini of SYP-2 and SYP-4 in *syp-4*^*CmutFlag*^ mutant animals in late pachytene. We were not able to map SYP-3 in *syp-4*^*CmutFlag*^ animals since *syp-3* and *syp-4* are closely linked on Chromosome I (Supplementary Table S1).

The results are summarized in Supplementary Table S2. All SYP-2, -3, and -4 epitopes localized near the midline of the SC in *x*, similar to “central element” proteins identified in other species (Fig. 6). We also observed consistent differences for most epitopes between early and late pachytene in wild-type worms and between wild-type and *syp-4*^*CmutFlag*^ animals. The C-terminus of SYP-2 localized to the center of the SC in *x* in early pachytene and *syp-4*^*CmutFlag*^ animals, but we were able to resolve a bimodal distribution with peaks at 18.7±2.4 nm from the center in wild-type animals in late pachytene (Fig. 6A). Similarly, the SYP-3 C-terminus was slightly more central in early pachytene (12.3±1.7 nm off center) than late pachytene (16.3±4.5 nm, Fig. 6B). By contrast, a GFP-tag at the N-terminus of SYP-3 localized to the SC midline in both early and late pachytene, suggesting a head-to-head arrangement of SYP-3 molecules (Fig. 6B). The thickness of the z-distributions of SYP-2 and SYP-3 epitopes increased from early to late pachytene in wild-type but not in *syp-4*^*CmutFlag*^ animals, similar to our findings for SYP-1 (Fig. 6A,B). Confocal microscopy further confirmed that the fluorescence intensity of *syp-4*^*CmutFlag*^ animals is lower than in control wild-type animals (Supplementary Fig. S5) further confirming that this mutant prevents or reduces the normal accumulation of SC proteins between homologs in late pachytene (Fig. 2C).

**Figure 6:**
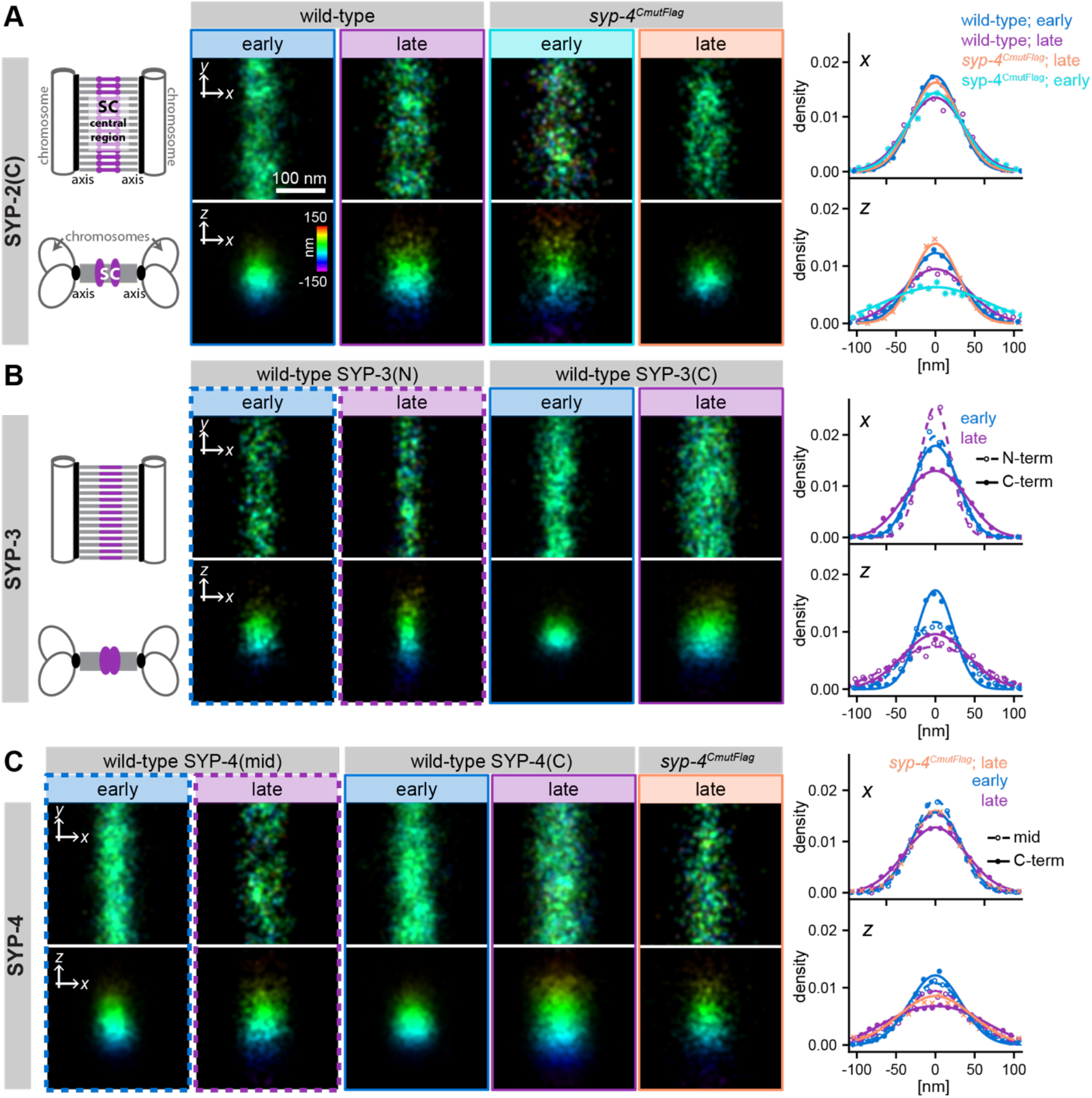
Individual components of the SC are distinctly reorganized during pachytene. (A) 3D-STORM images reveal opposing reorganizations of the C-terminus of SYP-2 during pachytene in wild-type and *syp-4*^*CmutFlag*^ animals in both, frontal (top) and cross-sectional (bottom) view. (B) Localizations of N- and C-terminus of SYP-3 in wild-type animals in early and late pachytene are shown. (C) Localizations of epitope tags inserted internally or at the C-terminus of SYP-4 in wild-type animals in early and late pachytene are shown. For comparison, the localization of the 3xFlag-tag in C-terminally mutated *syp-4*^*CmutFlag*^ animals is also shown. Scale bars in (A) apply to all images.

The differences in width along the *z*-axis in early vs. late pachytene were even more pronounced for the C-terminus of SYP-4: while it was confined to the central plane in early pachytene and *syp-4*^*CmutFlag*^ animals, we resolved a bimodal distribution along the *z* axis in wild-type late pachytene (Fig. 6C). Taken together, our data reveal striking changes within the ultrastructure of the SC during pachytene and in *syp-4*^*CmutFlag*^ mutant animals.

## Discussion

Through super-resolution imaging, we have found that SCs undergo a structural transition during pachytene. While some of our data can be explained by the accumulation of proteins within the SC, we also see evidence of reorientation of components relative to each other (Fig. 7). Moreover, we observed clear differences in the ultrastructure of both polycomplexes and SCs in *syp-4*^*CmutFlag*^ mutants. This aberrant SC ultrastructure is accompanied by an increase in the number of “designated crossovers” marked with GFP::COSA-1 foci, and a corresponding increase in genetically-detectable crossovers. While previous studies have revealed that the integrity of the SC is important for crossover interference (18, 20), (9, 14-18), (24), France:2021kb}, our findings demonstrate that continuous SC assembly along chromosomes is not sufficient for interference and support a role for SC ultrastructure in regulating CO distribution. However, how the SC mediates crossover interference is still contentious(71): Recent findings revealing that regulatory factors can diffuse along or within the SC (33, 72) suggested that interference between early recombination intermediates is mediated by a coarsening process driven by diffusion of pro-crossover factors along or within the SC (29, 31, 32, 73).

**Figure 7:**
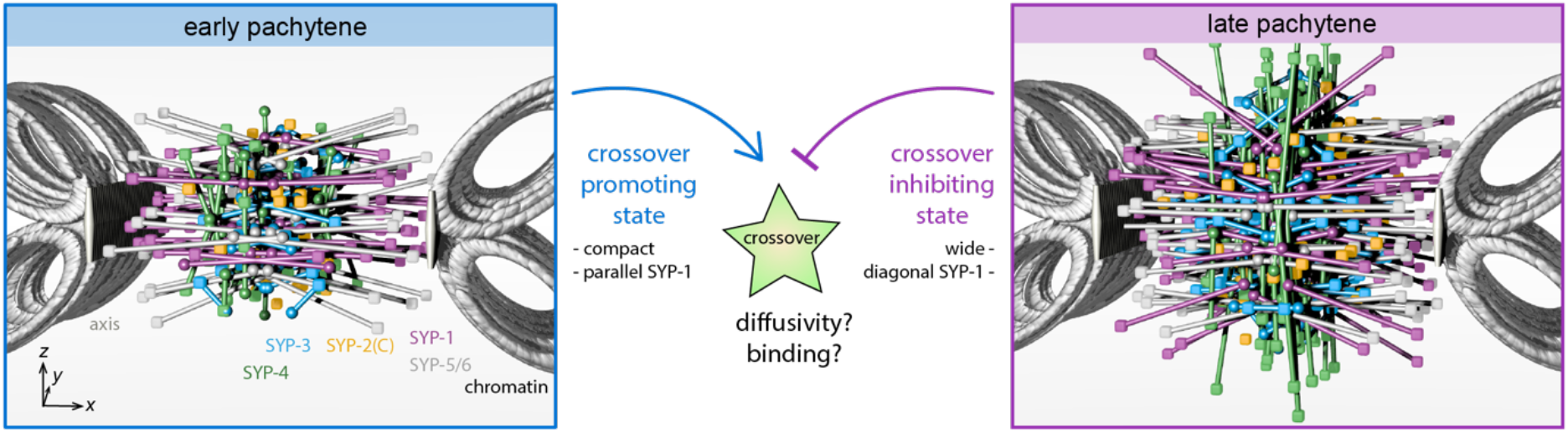
A probabilistic model reveals distinct ultrastructural states of the synaptonemal complex. The models of SC organization generated by mapping of N- and C-terminal distributions of SYP proteins is shown for early (left) and late (right) pachytene in wild-type animals. C-termini are represented as cubes and N-termini (central domain for SYP-4) as spheres. The localization of SYP-5/6 was mapped previously (18) and is shown for reference in gray. Interestingly, while SYP-1 are more splayed in late pachytene, SYP-5/6 are more parallel to the plane of the SC. The localization of HIM-3 in *x* is used as a proxy for axis localization. Gray loops represent sister chromatids of homologous chromosomes. Models are rendered using POV-ray (v3.7.0).

Our fortuitous isolation of a separation-of-function allele of *syp-4* reveals that defects in the ultrastructure of the SC are associated with severe defects in crossover interference: Changes in the SC ultrastructure may modulate the binding or diffusion of CO regulatory factors within the SC. Interestingly, perturbations applied to nematic liquid crystalline materials such as the SC(5) can change the orientation of molecules within the liquid crystal(74). Therefore, the designation of a CO event within the SC, which has been shown to distort the SC (30), may even trigger the reorganization of the SC we observed during pachytene in wild-type animals resulting in changes of the physical properties that may modulate the binding or diffusion of CO factors to establish interference along chromosomes (Fig. 7).

The C-terminal domain of SYP-4 is unique among *C. elegans* SC proteins: it lacks sequences predictive of coiled-coil structure, is likely to be intrinsically disordered, and has a very unusual amino acid composition. It is acidic and rich in glycine, which would make it very flexible. Most distinctively, it contains repeated instances in which phenylalanine alternates with other residues, most often asparagine, to form (F-x-F-x-F) motifs, and a terminal FxFF motif. These features are conserved within nematodes, which facilitated our identification of a SYP-4 ortholog in *Pristionchus pacificus*, a distant relative of *C. elegans* (75). Moreover, the C-terminus of SIX6OS1, an essential component of the central element in mice (76), contains very similar motifs, along with a similar amino acid composition and predicted disorder. An amino acid polymorphism in this domain was linked to elevated recombination rates in women in an Icelandic population (77). Thus, it seems likely that this C-terminal domain is a unique interface involved in crossover regulation, and that this function may be conserved between nematodes and mammals.

## Data Availability

NGS datasets generated in this study are available on the NCBI SRA database under accession number SRP126693 (https://www.ncbi.nlm.nih.gov/sra/SRP126693). Other data are available in the supplementary material (soure_data.xlsx).

## Acknowledgements

Some *C. elegans* strains used in this work were provided by the Caenorhabditis Genetics Center, which is funded by the NIH - Office of Research Infrastructure Programs (P40 OD010440). This work was supported by a postdoctoral fellowship of the Human Frontier Science Program to SK (LT000903/2013-C), an NSF Graduate Research Fellowship (DGE 1106400) to MW, the Pew Charitable Trusts and the Packard Fellowship for Science and Engineering to KX, and support to AFD from the National Institutes of Health (R01 GM065591) and the Howard Hughes Medical Institute.

## Notes

### Competing Interest Statement

The authors have declared no competing interest.

### Summary of Updates

The manuscript has been revised completely for clarity.

