## Supplementary Material for "Dynamic molecular architecture of the synaptonemal complex"

**Table S1: List of Strains used in this study and their characterization.** *p*-Values for deviation from wild-type are calculated using Mann-Whitney tests.

| name | genotype | egg viability [%] (*p*-value) | males [%] (*p*-value) | # of eggs |
|---|---|---|---|---|
| **N2** | *wild-type* | 104±6 | 0.08±0.20 | 4773 |
| **GFP::COSA-1** (60) | *meIs8* II | 107±4 | 0.00±0.00 | 1362 |
| **HA::SYP-1** | *meIs8* II; *mEos::him-3(ie33)* IV; *syp-1(ie40[syp-1(1-47)::HA::syp-1(48-489)])* V | 100±4 (0.886) | 0.00±0.00 (0.452) | 840 |
| **SYP-2::HA** | *meIs8* II; *mEos::him-3(ie33)* IV; *syp-2(ie99[syp-2::HA])* V | 101±6 (0.394) | 0.00±0.00 (1.00) | 1405 |
| **GFP::SYP-3** | *syp-3(ok758)* I; *ieSi11 [syp-3p::syp-3::EmGFP::syp-3-3'UTR + unc-119(+)]* II; *mEos::him-3(ie33)* IV | 83±10 (0.031) | 0.87±0.59 (0.045) | 1143 |
| **SYP-3::HA** | *syp-3(ie42[syp-3::HA])* I; *meIs8* II; *mEos::him-3(ie33)* IV | 95±4 (0.041) | 1.80±0.60 (0.004) | 1457 |
| **SYP-4$^{ha}$** | *syp-4(ie29[syp-4::HA])* I; *meIs8* II; *mEos::him-3(ie34)* IV | 100±2 (0.150) | 0.72±0.41 (0.052) | 1336 |
| **SYP-4$^{ha}$** | *syp-4(ie29[syp-4::HA])* I; *meIs8* II | 105±3 (0.686) | 0.00±0.00 (1.000) | 951 |

| | | | | |
|---|---|---|---|---|
| **SYP-4::intFlag** | *syp-4(ie30[syp-4(1-315)::2xFlag::syp-4(319-605)])* I; *meIs8* II; *mEos::him-3(ie33)* IV | 98±3 (0.032) | 1.91±0.59 (0.010) | 1350 |
| **SYP-4$^{CmutFlag}$** | *syp-4(ie25[syp-4ΔC::3xFlag])* I; *meIs8* II | 38±4 (0.008) | 5.97±1.25 (0.010) | 1024 |
| **SYP-4$^{3'UTR}$** | *syp-4(ie29[syp-4::HA::mod3'UTR])* I; *meIs8* II | 80±9 (0.002) | 1.58±1.58 (0.010) | 992 |
| **Δ*htp-3*** (5) | *htp-3(tm3655)* I; *meIs8* II | n.d. | n.d. | |
| ***Δhtp-3 syp-4$^{CmutFlag}$*** | *htp-3(ie100) syp-4(ie25)* I; *meIs8* II | n.d. | n.d. | |
| ***Δhtp-3 syp-4$^{3'UTR}$*** | *htp-3(ie101) syp-4(ie27)* I; *meIs8* II | n.d. | n.d. | |
| **SYP-4$^{CmutFlag}$** | *syp-4(ie25[syp-4ΔC::3xFlag])* I; *meIs8* II; *mEos::him-3(ie33)* IV | n.d. | n.d. | |
| **SYP-4$^{CmutFlag}$; HA::SYP-1** | *syp-4(ie25)* I; *meIs8* II; *mEos::him-3(ie33)* IV; *syp-1(ie40)* V | n.d. | n.d. | |

**Table S2: Summary of localizations of SC components determined by STORM.** Positions are distances of the maxima to the center of the SC (wt = wild-type; early = early pachytene; late = late pachytene). Position corresponds to the distance from the center of the SC for bimodal distributions, and HWHM denotes the half width at half maximum of the distributions.

| domain | condition | x [nm] | | z [nm] | | total length analyzed [nm] (# images) |
|---|---|---|---|---|---|---|
| | | position | HWHM | position | HWHM | |
| HA::SYP-1 | wt, early | 0 ± 0 | 28.4 ± 1.7 | 0 ± 0 | 33.5 ± 2.7 | 3438 (8) |
| | wt, late pachytene | 0 ± 0 | 26.9 ± 1.6 | 0 ± 0 | 29.7 ± 3.1 | 4780 (9) |
| | *syp-4*$^{CmutFlag}$; early | 0 ± 0 | 22 ± 1.3 | 0 ± 0 | 28.8 ± 0.7 | 4351 (10) |
| | *syp-4*$^{CmutFlag}$; late | 0 ± 0 | 20 ± 0.7 | 0 ± 0 | 20.1 ± 0.6 | 3114 (6) |
| SYP-1(C) | wt, early | 42.3 ± 0.9 | 22.8 ± 0.9 | 0 ± 0 | 38.4 ± 3.3 | 4092 (7) |
| | wt, late pachytene | 42.2 ± 1.2 | 28.4 ± 1 | 0 ± 0 | 65 ± 4.5 | 5312 (9) |
| | *syp-4*$^{CmutFlag}$; early | 38 ± 2.7 | 24.9 ± 0.8 | 0 ± 0 | 41.2 ± 5 | 2768 (5) |
| | *syp-4*$^{CmutFlag}$; late | 30.5 ± 1.4 | 29.1 ± 1.6 | 0 ± 0 | 53.9 ± 2.3 | 4848 (6) |
| SYP-2(C) | wt, early | 0 ± 0 | 27.2 ± 1.8 | 0 ± 0 | 37.7 ± 3.5 | 5866 (10) |
| | wt, late pachytene | 18.7 ± 2.4 | 21.8 ± 1.5 | 0 ± 0 | 49.9 ± 6.8 | 4573 (9) |
| | wt, late (HA) | 20.8 ± 0.9 | 20 ± 1.1 | 0 ± 0 | 37.6 ± 3.1 | 2958 (8) |
| | *syp-4*$^{CmutFlag}$; early | 0 ± 0 | 32.2 ± 2.1 | 0 ± 0 | 80.5 ± 7.4 | 3172 (7) |
| | *syp-4*$^{CmutFlag}$; late | 0 ± 0 | 29 ± 2.5 | 0 ± 0 | 32 ± 2.6 | 3604 (8) |
| GFP::SYP-3 | wt, early | 0 ± 0 | 23.5 ± 0.6 | 0 ± 0 | 41.4 ± 3.3 | 1784 (4) |
| | wt, late pachytene | 0 ± 0 | 18.2 ± 1.3 | 0 ± 0 | 59.9 ± 5.5 | 2238 (6) |
| SYP-3::HA | wt, early | 12.3 ± 1.7 | 20.3 ± 1.1 | 0 ± 0 | 27.4 ± 2.1 | 4653 (10) |
| | wt, late pachytene | 16.3 ± 4.5 | 28.6 ± 2.2 | 0 ± 0 | 49.6 ± 6.3 | 2764 (7) |

| | | | | | | |
|---|---|---|---|---|---|---|
| **SYP-4::intFlag** | wt, early | 0 ± 0 | 25.8 ± 0.8 | 0 ± 0 | 47 ± 4 | 2981 (8) |
| | wt, late pachytene | 0 ± 0 | 30.2 ± 3.4 | 0 ± 0 | 49.7 ± 3.7 | 2524 (4) |
| **SYP-4::HA** | wt, early | 14.2 ± 2.1 | 22.9 ± 1.5 | 0 ± 0 | 40.5 ± 2.1 | 5355 (10) |
| | wt, late pachytene | 0 ± 0 | 36.8 ± 1.2 | 30.6 ± 11.4 | 51 ± 8.7 | 5264 (9) |
| **SYP-4$^{CmutFlag}$** | *syp-4$^{CmutFlag}$*; late | 0 ± 0 | 29.2 ± 1.2 | 0 ± 0 | 56.7 ± 5.4 | 4611 (9) |
| **HIM-3** | wt, early | 33.1 ± 4.7 | 43.3 ± 2.1 | n.d. | n.d. | 2548 (6) |
| | wt, late pachytene | 47.3 ± 1.7 | 30.1 ± 1.3 | 0 ± 0 | 31.4 ± 2.9 | 4411 (11) |
| | *syp-4$^{CmutFlag}$*; early | 46.1 ± 3.8 | 48.4 ± 2.9 | n.d. | n.d. | 3900 (7) |
| | *syp-4$^{CmutFlag}$*; late | 36 ± 4 | 45.9 ± 3.3 | n.d. | n.d. | 2926 (6) |

**Table S3: List of synthetic oligos and gRNAs**

| Name | Description/Target | Sequence |
|---|---|---|
| crSK3 | crRNA: syp-1 (N-terminus) | CGAGCAAACTGATGAGCAGG |
| crSK7 | crRNA: syp-4 (internal) | AATTCCGACACAGCATCCGA |
| crSK9 | crRNA: syp-2 (C-terminus) | TGGTTGAAACGCTCGAGCCG |
| crSK10 | crRNA: syp-3 (C-terminus) | TTTAATTCATGTAGAAAGTC |
| crSK14 | crRNA: syp-4 (C-terminus) | GGAGCAACTTCTGGAGCGGG |
| SK251 | repair template: syp-4::ha | GAGATGGATCGTTCAACTTTAACTTTGACGGTGACGGTGAAGGCGGAGCAACTTCaGGcGCtGGCGGAAACAGCACCTCGTTCTTTAACTTTTACCCCTACGATGTCCCAGATTATGCTTAGaaaattatcatgtattattcagctcttgtatcatttgtatcg |
| SK189 | repair template: syp-4::3xFlag(C) | TAACTTTGACGGTGACGGTGAAGGCGGAGCAACTTCTGGcGCtGGCGGAAACAGCACCTCGTTCTTTAACTTTGACTATAAAGATCACGACGGAGATTACAAGGACCATGATATCGACTACAAGGACGACGACGACAAGGGATAGaaaattatcatgtattattcagctcttgtatcatttgtatcgtttaatgtcagaa |
| SK250 | repair template: syp-4::Flag(internal) | GCGAAACCGCGAAATCTTCCTTCTCCAAAGAGCCAATTCCGACtCAaCATGACTATAAAGATCACGACGGAGATTACAAGGACCATGATATCGACTACAAGGACGACGACGACAAGGGACCaATGGAAGTTGGCGAAGAACCTGTAATCGAACATTCAGCTGAAGAAAGCG |
| SK285 | repair template: ha::syp-1 | gagATCATGCTCACACAGCAATACGTTCTTGAGAAGCTTGACGAGCAGACTTACCCCTACGATGTCCCAGATTATGCTGAGCAAGATAAAGGCGAGCAAACTGATGAGCAGGCtGCGAAATCTCGCGAGCTGGCTTCTCAAGTTCAAACC |
| SK302 | repair template: syp-2::ha | GGAAGATTTCTCTGCTCATTACGACAAACTTCTGGATTTGGTTGAgACcCTtGAaCCaTGGGCTGACAAGTTATACCCCTACGATGTCCCAGATTATGCTTAAatcatctgttgtattcaatttccttgttttattacatgacatg |
| SK303 | repair template: syp-3::ha | GAGTACCAAAAATGTCGTCTACAAGAAGAAGCTCTGAAAGCtCGACTTTCTACATACCCCTACGATGTCCCAGATTATGCTTGAattaaaagaatcagaacagccaaaatgtgggcaactgttaatcgaactgtaatg |

| | | |
|---|---|---|
| SK189 | genotyping primer, forward: syp-4 (C-terminus) | GGATGAAGGAAAGGGTTCGGG |
| OR236 | genotyping primer, reverse: syp-4 (C-terminus) | GAGGCAATTTCTTGGTCCAATAA |
| SK239 | genotyping primer, forward: syp-4 (internal) | GTTTGGAAACATTGAAGGACGAGG |
| SK142 | genotyping primer, reverse: Flag in syp-4 (internal) | TCGTCCTTGTAGTCGATATCATGGT |
| SK199 | genotyping primer, forward: syp-1 (N-terminus) | acgcgctccattgacaaaatc |
| SK201 | genotyping primer, reverse: syp-1 (N-terminus) | TCATGTTTCTGCTTGGCTGC |
| SK220 | genotyping primer, forward: syp-2 (C-terminus) | GAGCTCGTCGAGAAAGCCAAC |
| SK221 | genotyping primer, reverse: syp-2 (C-terminus) | cctgtgtcctatgcggcg |
| SK304 | genotyping primer, forward: syp-3 (C-terminus) | GCAGAGAAACAACTAGAAGTCGCC |
| SK305 | genotyping primer, reverse: syp-3 (C-terminus) | aatttattaaaacttcacaacggttgatgc |

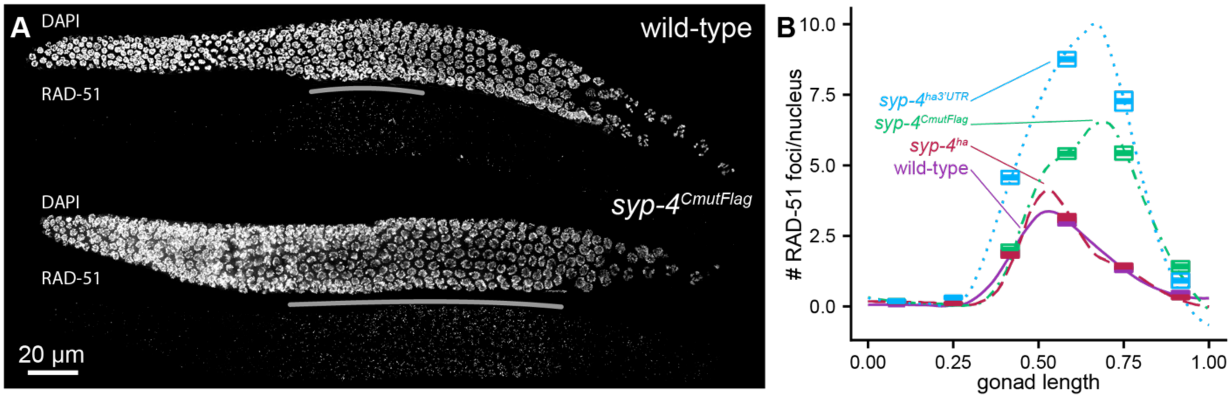


**Figure S1: Double-strand break formation is extended during *syp-4*$^{CmutFlag}$ meiosis.** (A) Projection images show DAPI (upper) and RAD-51 immunofluorescence (lower), which marks double-strand breaks. While the RAD-51 zone (gray line) is normally limited to a short region in early/mid-pachytene in wild-type (top), it is markedly elongated in *syp-4*$^{CmutFlag}$ (bottom). (B) shows the average number of RAD-51 foci per nucleus (boxes are mean±s.e.) as a function of the relative length of the gonad from the distal tip to the end of pachytene of wild-type (purple solid line, n=1672 nuclei from (7) gonads), *syp-4*$^{ha}$ (red dashed line, n=1407 (6)), *syp-4*$^{CmutFlag}$ (green dot-dashed line, n=1440 (6)) and *syp-4*$^{ha3'UTR}$ (blue dotted line, n=1162 (4)), respectively.

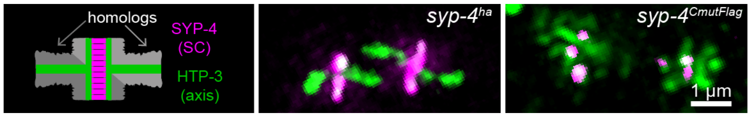


**Figure S2: Bivalent structure is perturbed in *syp-4$^{CmutFlag}$* animals**. The cartoon (left) illustrates the normal cruciform bivalent structure stained by HTP-3 (green), while SYP-4 (magenta) is restricted to the short arm. While *syp-4$^{ha}$* animals (center) show typical cruciform structures, the bivalents are disrupted in *syp-4$^{CmutFlag}$* (right), likely reflecting the occurrence of multiple crossovers per chromosome pair.

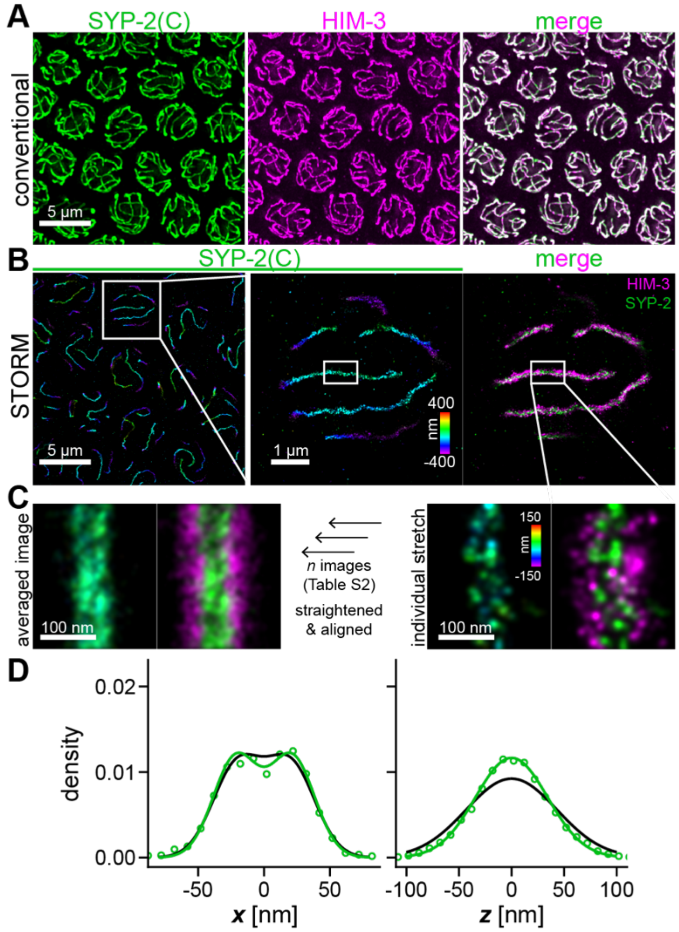


**Figure S3: Experimental strategy for mapping the architecture of the synaptonemal complex.** (A) Diffraction-limited fluorescence microscopy cannot resolve the internal organization of the axis or SC (HIM-3, magenta, left) and SC components (SYP-2::HA, green, center) are co-localized (merge, right). (B) The structure of the SC is resolved by STORM super-resolution microscopy. Colors denote localization in *z* (left and center). Localization of GFP::HIM-3 (magenta, right) is used to determine the orientation of the SC (SYP-2::HA(C), wild-type, late pachytene in green). White boxes in B indicate the regions enlarged in other panels. (C) For further analysis, individual stretches in frontal view (right), which are characterized by clearly separated HIM-3 axes, are rotated, straightened, and aligned in *x* and *z* to generate the averaged image (left). (D) The results are highly reproducible between different preparations and different antibodies (SYP-2(C) peptide antibody, black, and SYP-2::HA epitope antibody, green) (for results of fit see Table S2). Note that the measured distribution is narrower for a monoclonal anti-HA antibody than for a polyclonal anti-SYP-2(C) peptide antibody.

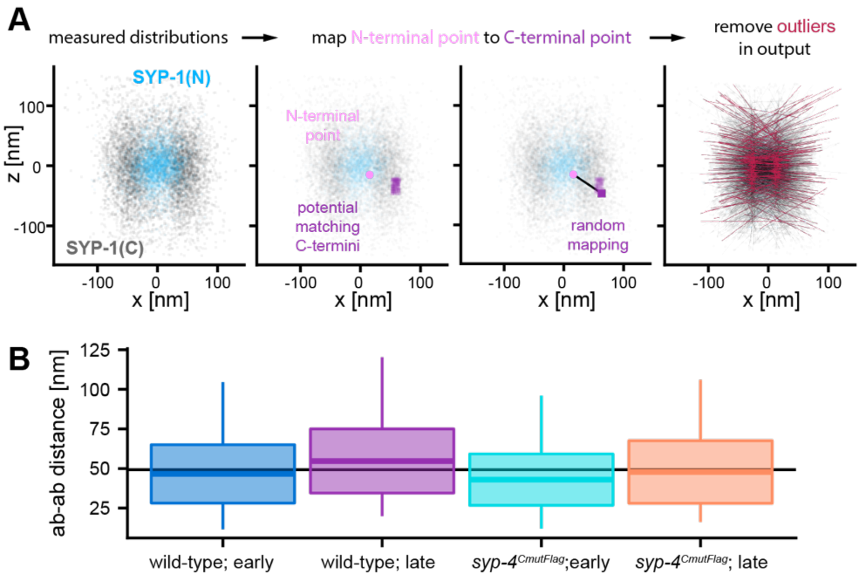


**Figure S4: Dynamic orientation of SYP-1 molecules.** (A) To generate a model of SYP-1 orientation within the SC, individual datapoints (magenta circle, center) in the measured distributions of the N- (blue, left) and C-terminus (gray, left) of a given component (here: SYP-1 in wild-type late pachytene) were mapped to a random localization event within the same percentile ±7.5% tolerance of the C-terminal distribution in $x$ and $z$ (purple, center). Mapped events resulting in the shortest or longest 5% of antibody-to-antibody distances (red, right) were disregarded. (B) The boxplots show means±s.d. of the distances between N- and C-terminal antibodies as estimated by this probabilistic mapping approach. Whiskers are extremes. The black line shows the mean across all conditions and genotypes (49.3±6.9 nm (s.d.)).

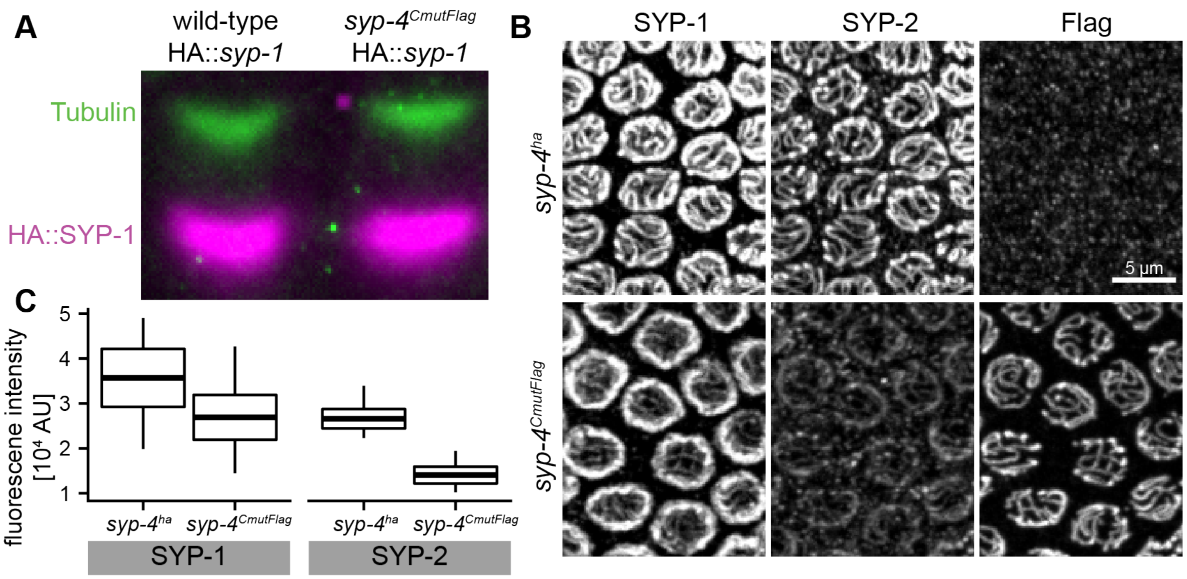


**Figure S5: Abundance of SC proteins in *syp-4$^{CmutFlag}$*.** (A) The expression level of HA::SYP-1 (magenta) is indistinguishable between wild-type and *syp-4$^{CmutFlag}$* strains, as shown by a western blot. Tubulin (green) is shown as a reference. (B) By contrast, immunofluorescence intensities for SYP-1 (left) and SYP-2 (center) along SCs are markedly reduced in *syp-4$^{CmutFlag}$* SCs (bottom) compared to wild-type (*syp-4$^{ha}$*, right). To ensure identical handling, dissected worms of both genotypes were mixed and stained together, and their genotypes were determined by anti-FLAG staining (right), which detects SYP-4 in *syp-4$^{CmutFlag}$* but not in *syp-4$^{ha}$* germlines. Note that this SYP-1 antibody also recognizes a nuclear pore protein, which becomes more apparent in *syp-4$^{CmutFlag}$* animals. (C) Boxplots show mean±s.d. (whiskers are extreme values) of average fluorescence intensities of late pachytene SCs (*syp-4$^{ha}$*: *n*=339, *syp-4$^{CmutFlag}$*: *n*=209). SCs were segmented using SYP-2 (*syp-4$^{ha}$*) or Flag (*syp-4$^{CmutFlag}$*) staining.
